# Ubiquitin- and ATP-dependent unfoldase activity of P97/VCP•NPLOC4•UFD1L1 is enhanced by a mutation that causes multisystem proteinopathy

**DOI:** 10.1101/129528

**Authors:** Emily E. Blythe, Kristine C. Olson, Vincent Chau, Raymond J. Deshaies

**Affiliations:** Division of Biology and Biological Engineering, California Institute of Technology, Pasadena CA, 91125; Pennsylvania State University College of Medicine, Hershey PA, 17033; Howard Hughes Medical Institute, California Institute of Technology, Pasadena CA, 91125

## Abstract

p97 is a ‘segregase’ that plays a key role in numerous ubiquitin-dependent pathways, such as ER-associated degradation (ERAD). It has been hypothesized that p97 extracts proteins from membranes or macromolecular complexes to enable their proteasomal degradation; however, the complex nature of p97 substrates has made it difficult to directly observe the fundamental basis for this activity. To address this issue, we developed a soluble p97 substrate—Ub-GFP modified with K48-linked ubiquitin chains—for in vitro p97 activity assays. We demonstrate for the first time that wild type p97 can unfold proteins and that this activity is dependent on the p97 adaptor NPLOC4-UFD1L, ATP hydrolysis, and substrate ubiquitination, with branched chains providing maximal stimulation. Furthermore, we show that a p97 mutant that causes inclusion body myopathy, Paget’s Disease of bone, and frontotemporal dementia (IBMPFD) in humans unfolds substrate faster, suggesting that excess activity may underlie pathogenesis. This work overcomes a significant barrier in the study of p97 and will allow the future dissection of p97 mechanism at a level of detail previously unattainable.

## Introduction

The AAA ATPase p97, also called valosin containing protein (VCP) or Cdc48 in yeast, is integral to a wide array of processes in the cell (1, 2). Its role in ER-associated degradation (ERAD) is the most well-studied, where it functions to pull misfolded proteins from the ER membrane so they can be degraded by the proteasome (3). Other degradative pathways with substrates embedded in large structures, such as ribosome associated degradation (RAD) and mitochondrial associated degradation (MAD), also rely on the activity of p97 (4–6). Furthermore, p97 is involved in non-degradative pathways like Golgi and nuclear envelope reassembly after mitosis and endosomal trafficking (7, 8). The common mechanism that underlies all of these cellular jobs is presumed to be the extraction and unfolding of ubiquitylated proteins by p97 (9, 10). Its homology to other AAA ATPases that have demonstrated unfolding activities, such as the proteasome 19S regulatory particle (11, 12), ClpA (13), and VAT (14, 15), further support the model of p97 as an unfoldase. While p97 has been shown to extract proteins from membranes and DNA, its ability to unfold a protein has not been explicitly demonstrated. Because of this considerable gap in our knowledge, the biochemical activity that underlies p97’s myriad functions remains a mystery.

Despite our poor molecular understanding of exactly what p97 does and how it does it, structural studies and indirect assays have provided some insight into p97 activity. p97 is a homohexamer, with each protomer comprising an N-domain, two ATPase domains (D1 and D2) stacked one upon the other, and an unstructured C-terminal tail (16–18). Whereas both ATPase domains are functional, it is thought that hydrolysis of ATP in D2 is the main driver of mechanical force and ATP binding in D1 promotes hexamer formation (19–25). The conformations of the two ATPase domains and the N-domain with respect to one another are highly cooperative and dependent upon nucleotide binding and hydrolysis (17, 18, 20, 22, 26–32). Recent high-resolution structural studies show movement of D2 relative to N-D1 upon ATP binding in D2, while the N domains move from a coplanar to an axial position with respect to D1 upon ATP binding to D1 (18). It is unclear how these conformational changes translate to mechanical force for remodeling protein substrates. Homologues like ClpA function by threading polypeptides through the central pore (33), and though p97 lacks the requisite hydrophobic pore residues in D1, other pore residues interact with and are key to the processing of substrates (19, 34). However, structural studies imply that the central pore of p97 may be too narrow to accommodate a polypeptide (16, 18). Other proposed mechanisms include unfolding by D2 through the arginine “denaturation collar” and unfolding outside of the pore by the movement of the N-domains similar to what has been proposed for NSF (34, 35). Resolution of the key question of how p97 works has been stymied by the lack of a direct assay to measure the core biochemical activity underlying its segregase function.

In performing its biological functions, p97 does not act alone but instead associates with a set of adaptor proteins that act to recruit or modify substrates (36). Many of the adaptors contain ubiquitin-binding domains or can add or remove ubiquitin modifications, thereby linking p97 to ubiquitin signaling (36, 37). Some adaptors also modulate p97 ATPase activity (38–40). Whereas a few adaptors have been linked to specific pathways or substrates (41, 8, 42–44), the functions of others remain unknown (36, 37). The best characterized p97 adaptor is the heterodimer of NPLOC4/Npl4 and UFD1L/Ufd1 (UN), which recruits substrates in ERAD and other proteasome-dependent degradative processes (45–47). NPLOC4 and UFD1L each bind ubiquitin chains, with UFD1L showing specificity for K48-linked chains (25, 48, 49). They interact with p97 at separate sites to form a complex with the stoichiometry of one UN heterodimer per p97 hexamer (39, 46, 50–52).

A significant incentive to gaining a deeper understanding of p97 mechanism of action is the deep connection between this protein and human disease, and possibly cancer therapy. Human p97 is mutated in the inherited, autosomal-dominant multisystem proteinopathy known as IBMPFD (53, 54). In addition, a small fraction of patients with inherited Amyotrophic Lateral Sclerosis (ALS/Lou Gehrig’s Disease) also carry mutations in p97 that overlap with those seen in IBMPFD patients (54, 55). However, the mechanism of pathogenesis is not understood in either case. It has been suggested at various times that pathogenesis arises from a failure of autophagy, endosomal sorting, clearance of leaky lysosomes, mitochondrial homeostasis, or mTOR regulation (56, 57, 8, 58– 61). Regardless of the cellular target, the molecular-level defect remains obscure. It has been suggested that failure of mutant p97 to bind UBXN6/UBXD1 is key (8), but IBMPFD mutants also show increased binding of UN (62). The mutations that cause IBMPFD all fall within the N-D1 domain interface and affect the relative orientation of these domains (63–66). Moreover, the mutations cause elevated ATP hydrolysis (20, 63–65, 67), but it has been suggested that this is an indirect consequence of a decoupling of substrate binding from mechanochemical transduction in the D2 domain (54, 68). Because p97 is a hexamer, it has been unclear how to interpret the autosomal dominant nature of IBMPFD. Is this truly due to enhanced activity, or do the mutations actually cause reduction-of-function through a dominant-negative mechanism, with mutant protomers poisoning mixed hexamers? Studies in Drosophila support the idea that the pathogenesis of IBMPFD mutations stems from elevated p97 activity, resulting in increased processing of TDP-43 (69) and mitofusin (61). As of yet there remains no biochemical assay that measures p97’s presumed core function of protein unfolding that would enable a direct test of this hypothesis in a defined system.

The nature of the majority of known p97 substrates—unstable, scarce, modified by ubiquitin, and not readily divorced from their contexts—presents challenges for studying the enzymatic activity of p97 in a systematic manner. A major barrier to progress has been the absence of a simple, rapid, quantitative assay using defined components, that can be employed to dissect in detail the mechanism of action of p97. To address this obstacle, we have developed a soluble, monomeric p97 substrate. Our substrate is based on a non-cleavable ubiquitin fusion protein, Ub^G^76^V^GFP, which is targeted for proteolysis by the Ub fusion degradation (UFD) pathway (70). Normally, ubiquitin fusions are co-translationally cleaved by a deubiquitinating enzyme to remove the ubiquitin (71). However, if the C-terminal glycine is mutated, processing is blocked and the fusion is rapidly degraded. Previous studies have demonstrated that the degradation of these non-cleavable Ub fusion proteins, including Ub^G^76^V^GFP, is dependent upon p97•UN in human, *Drosophila*, and yeast cells (19, 47, 70, 72). We show that p97 can unfold Ub^G^76^V^GFP modified with a K48-linked polyUb chain and that this reaction is dependent upon the nature of the Ub chain, UN, and p97 ATPase activity in D2. Our system provides the first direct demonstration of a p97 unfoldase activity that depends on predicted physiological requirements and will be an invaluable tool for further study of p97 mechanism.

## Results

### Substrate and assay design

We chose to pursue the UFD pathway substrate Ub^G^76^V^GFP because it is rapidly degraded in a p97-dependent manner in yeast, *Drosophila*, and human cells (19) and is a well-behaved protein whose folding state can be easily monitored by fluorescence. Since p97 substrates are often polyubiquitylated, and p97•UN binds polyubiquitin (25), we reasoned that Ub^G^76^V^GFP would need to be polyubiquitylated to be recognized. To efficiently ubiquitylate it, we developed a chimera of the RING domain from the E3 ubiquitin ligase gp78 and the E2 enzyme Ube2g2. Prior studies have shown that these enzymes catalyze formation of K48-linked ubiquitin chains (25). Notably, these two enzymes function upstream of p97 in ERAD, attesting to the physiological relevance of using these enzymes to generate a p97 substrate for our assay (73, 74). As compared to Ube2g2 alone or unfused Ube2g2 with added gp78RING (Fig. S1*A*), the gp78RING-Ube2g2 chimera produced unanchored polyUb chains with very high efficiency (Fig. S1*B*).

Using this tool, we employed various strategies to produce three types of potential p97 substrates. To simplify notation, linearly fused proteins are shown by dashes, and the length of the K48-linked Ub chain attached to a particular Ub is shown with a superscript preceding the initiator Ub. First, we aimed to create a substrate with a short Ub chain of defined length. Although the minimal requirement for recognition of ubiquitylated proteins by p97 is not known, the minimum Ub chain length for recognition by the proteasome is four (75). Therefore, we enzymatically ligated Ub_3_, in which the ubiquitins were joined via K48 linkages and the distal ubiquitin carried a K48R mutation, onto the linearly fused Ub to form pure ^Ub^3^^Ub-GFP (Fig.1A) (76, 77). Second, to create a substrate with longer polyubiquitin chains, we built K48-linked chains directly onto the linearly fused Ub (Fig. 1B). Finally, to produce substrate with branched Ub chains, we built Ub chains on a base substrate containing two or more linearly-fused Ub (Ub-Ub-GFP or Ub-Ub-Ub-GFP) (Fig. 1C). Heterogeneous substrates were fractionated by size exclusion chromatography to enrich for different chain lengths (Fig. 1D).

**Fig. 1.**
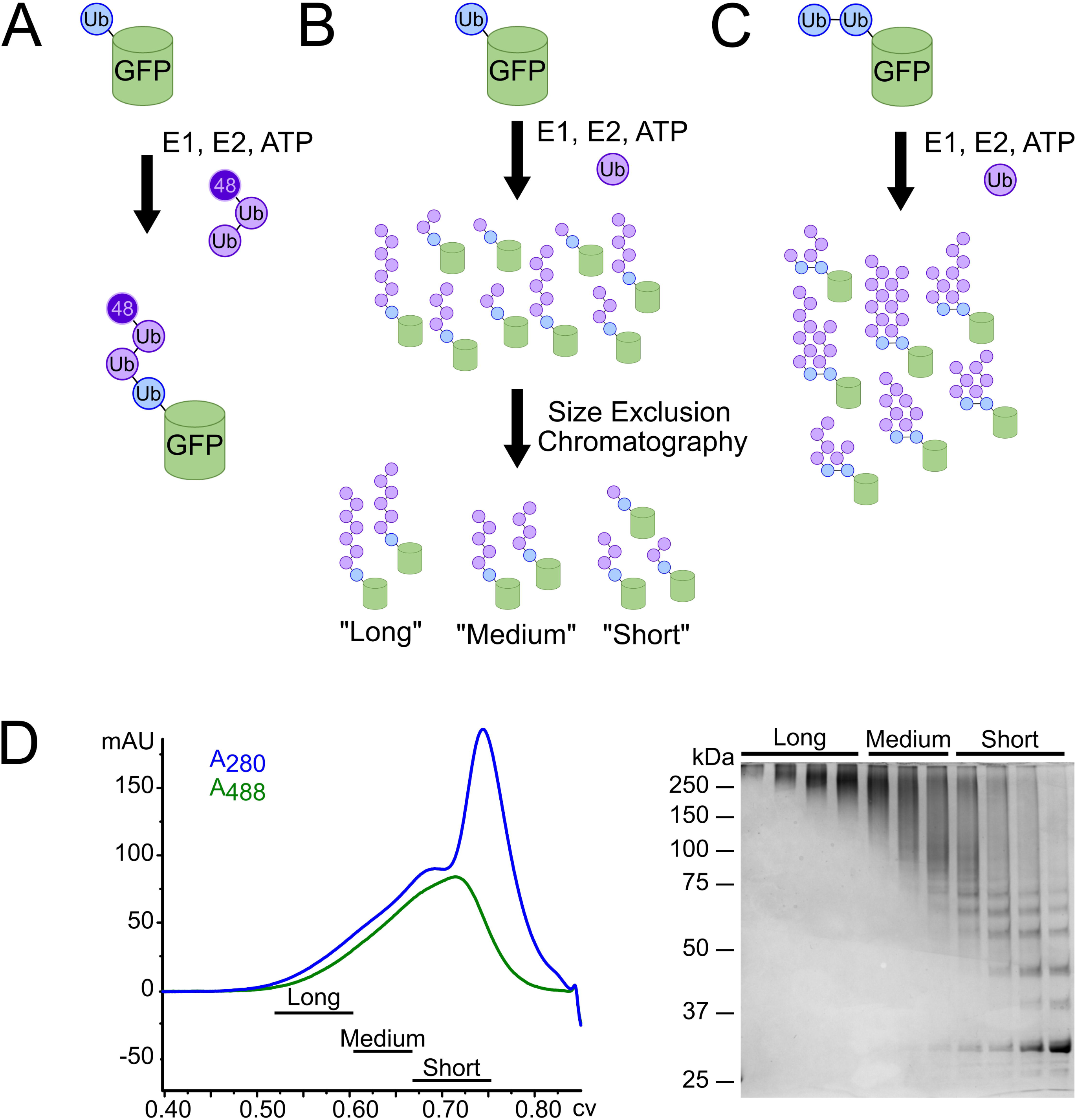
Substrate design and synthesis. (*A*) Preassembled, K48-linked Ub_3_ chains containing a K48R mutation on the distal ubiquitin were ligated onto a non-cleavable linear His_6_-Ub-GFP fusion protein to produce pure ^Ub^3^^Ub-GFP. (*B*) E1, E2, ubiquitin, and ATP were added to His_6_-Ub-GFP to elongate K48-linked ubiquitin chains of varying length on the ubiquitin fused to GFP. These resulting substrates were purified from free ubiquitin chains via Ni-NTA resin and crudely fractionated according to chain length via size exclusion chromatography to produce pools of “long,” “medium,” and “short” chain substrates (^UbL^Ub-GFP, ^UbM^Ub-GFP, and ^UbS^Ub-GFP, respectively). (*C*) To produce branched chains, Ub chains of varying length were enzymatically elongated on a di-or tri-ubiquitin linear fusion protein, Ub-Ub-GFP or Ub-Ub-Ub-GFP, similar to (*B*). (*D*) Size exclusion chromatogram and corresponding SDS-PAGE gel for the purification of substrate described in (*B*).

One concern we had was the potential for GFP to refold after being processed by p97, leaving the assay without an observable endpoint. To address this, we added an ATPase mutant of the chaperonin GroEL. The GroEL D87K ‘trap’ retains the ability to bind unfolded proteins but can no longer release those proteins (78). This dead-end complex sequesters unfolded GFP, preventing it from refolding, and has been used previously to provide assay endpoints for other unfolding machines (13).

### GFP is unfolded by p97 in an Ub- and UN-dependent manner

To explore the unfolding potential of p97, we first compared a set of Ub-GFP substrates bearing K48-linked ubiquitin chains of varying lengths (Fig. 2A). When mixed with p97, UN, and GroEL (Fig. S2), Ub-GFP and ^Ub^3^^Ub-GFP showed no appreciable loss of GFP fluorescence (Fig. 2B). However, Ub-GFP with both “medium” (>4 Ub, ^UbM^Ub-GFP) and “long” (>12 Ub, ^UbL^Ub-GFP) ubiquitin chains showed a modest decrease in fluorescence over time (∼30%, Fig. 2B). Whereas ^UbL^Ub-GFP did show improved unfolding relative to ^UbM^Ub-GFP, the difference was quite small (∼5% signal loss).

**Fig. 2.**
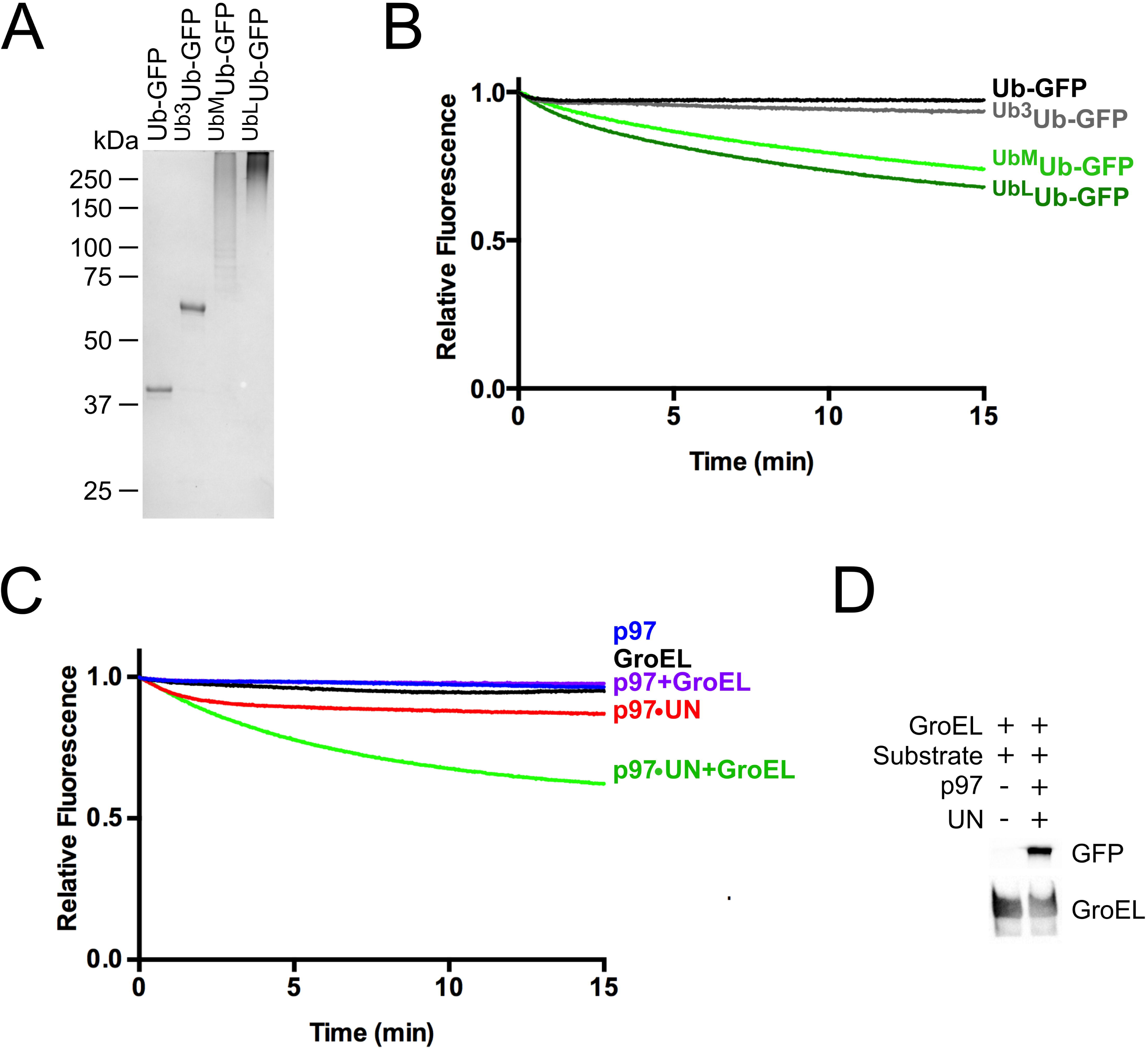
p97 unfolds Ub-GFP in a UN-dependent manner. (*A*) SDS-PAGE analysis of GFP substrates with different Ub chain structures stained with Coomassie. (*B*) Upon addition of ATP, 75 nM p97, 150 nM UN, and 250 nM GroEL, 25 nM Ub-GFP and ^Ub^3^^Ub-GFP did not appreciably lose fluorescence over time. However, Ub-GFP with “medium” or “long” K48-linked chains (^UbM^Ub- GFP and ^UbL^Ub-GFP) exhibited 26% and 32% loss of signal after 15 minutes, respectively. Representative traces shown, n≥3. (*C*) Fluorescence of ^UbL^Ub-GFP did not change over time with the addition of p97, GroEL, or p97 plus GroEL. Upon addition of p97 plus UN, a small decrease in signal was observed, and this decrease was augmented with the addition of GroEL. Representative traces shown, n≥2. (*D*) ^UbL^Ub-GFP co-immunoprecipitated with GroEL only in the presence of p97 and UN.

We examined further the requirements for p97-dependent unfolding using ^UbL^Ub- GFP, since this substrate gave the largest signal. When incubated with only p97 or p97 and GroEL, no unfolding was observed (Fig. 2C). A similar result was obtained when UN was replaced with NSFL1C/p47 or UBXN7/UBXD7, p97 adaptors involved in Golgi reassembly after mitosis (41) and regulation of cullin-RING Ub ligases (37, 43, 79), respectively, despite both of these adaptors binding to p97 and substrate (Fig. S3). Therefore, UN is required for p97- catalyzed unfolding, and it cannot be replaced by other adaptors. Additionally, GroEL was essential to provide an endpoint for the assay. Fluorescence loss was amplified by the addition of GroEL (Fig. 2C), indicating GFP was able to refold to some degree after processing by p97•UN. However, GroEL did not unfold substrate on its own (Fig. 2C), and immunoprecipitation of GroEL showed that p97•UN was required for GroEL interaction with substrate (Fig. 2D). Unfolding of substrate by p97 was also highly temperature dependent, with the rate and extent of unfolding increasing between 22-42 °C (Fig. S4).

We found it curious that ≤40% of the fluorescence signal of ^UbL^Ub-GFP was typically lost in our unfolding assays even though all components of the system were at or very near saturation (Fig. S5), suggesting that there was another factor influencing substrate competence that remained to be discovered. Three Ub binding sites with different chain linkage preferences are available on p97•UN (25). Therefore, we tested whether substrates carrying branched Ub chains would be more effectively unfolded, because a branch would enable two separate ubiquitin chains to be elaborated from a single attachment point (in this case, Met1 of GFP). As a proxy for Ub chains with branched linkages, we expressed Ub-GFP fused to one or more additional ubiquitins in tandem and used these proteins as substrates for subsequent enzymatic polyubiquitylation (Fig. 3A). Interestingly, a substrate in which K48-linked polyubiquitin chains were polymerized on Ub-Ub-GFP (^UbM^Ub-^UbM^Ub-GFP) showed significant improvement in unfolding by p97 as compared to ^UbM^Ub-GFP (Fig. 3B) despite the latter having a greater amount of ubiquitin conjugation as judged by mobility upon SDS-PAGE (Fig. 3A, lanes 1 and 2). Neither further extension of the chains to form ^UbL^Ub- ^UbL^Ub-GFP nor use of a triubiquitin fusion significantly increased the rate or extent of unfolding (Fig. 3B). Although we cannot directly visualize ubiquitin chains branching from each ubiquitin in the Ub-Ub-GFP fusion protein, reactions run with Ub-K48R indicate that both Ub moieties were efficiently conjugated with ubiquitin under our reaction conditions (Fig. S6). Incidentally, this same reaction confirms the linkage specificity of our Ube2g2–gp78 E2–E3 chimera. Taken together, our data suggest that the physical arrangement of the Ub chains is important for unfolding by p97•UN, with the enzyme preferring substrates with at least one branch point that enables nucleation of more than one chain of K48- linked ubiquitins.

**Fig. 3.**
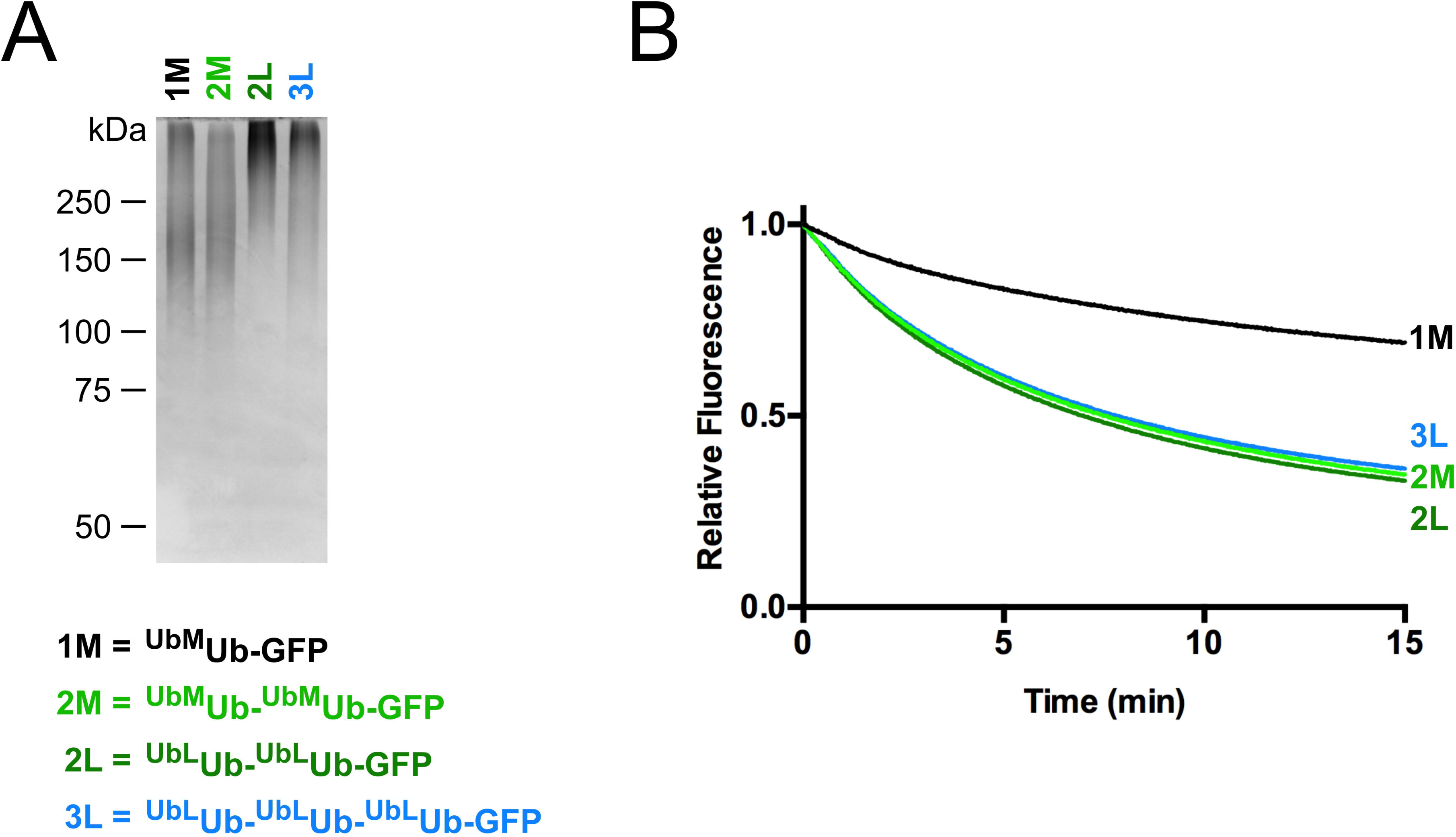
Branched Ub chains are better p97 substrates. (*A*) SDS-PAGE analysis of Ub-GFP substrates stained with Coomassie. (*B*) Comparison of unfolding of substrates with one, two or three ubiquitins fused in tandem to GFP. Note that substrate with two linearly fused ubiquitins (e.g. ^UbM^Ub-^UbM^Ub-GFP) was unfolded to a greater extent by p97 than substrate with a single linearly-fused ubiquitin, even though aggregate ubiquitination for the latter substrate was at least as extensive or greater than the former. Adding an additional linearly-fused ubiquitin (^UbL^Ub-^UbL^Ub-^UbL^Ub–GFP) yielded no further improvement. Representative traces shown, n≥3.

### Substrate unfolding is dependent upon ATP hydrolysis and stimulates p97 ATPase activity

Next we examined the energy-dependence of p97-catalyzed unfolding. ^UbL^Ub- ^UbL^Ub-GFP was not unfolded by p97 in the absence of nucleotide, and ADP or the nonhydrolyzable ATP analog ATP?S could not substitute for ATP (Fig. 4A). Two p97 ATPase inhibitors, the allosteric inhibitor NMS-873 (80) and the D2-specific, ATP-competitive inhibitor CB-5083 (81), also prevented ATP-dependent substrate processing (Fig. 4A). p97 with a D1 domain Walker B motif mutation (p97-E305Q) that blocks nucleotide hydrolysis but not binding exhibited only mild defects in substrate unfolding. By contrast, the same mutation in D2 (p97- E578Q) completely abolished unfoldase activity (Fig. 4B; Table 1). Together, these data demonstrate that ATP hydrolysis in D2 powers the unfolding of substrate by p97•UN.

**Fig. 4.**
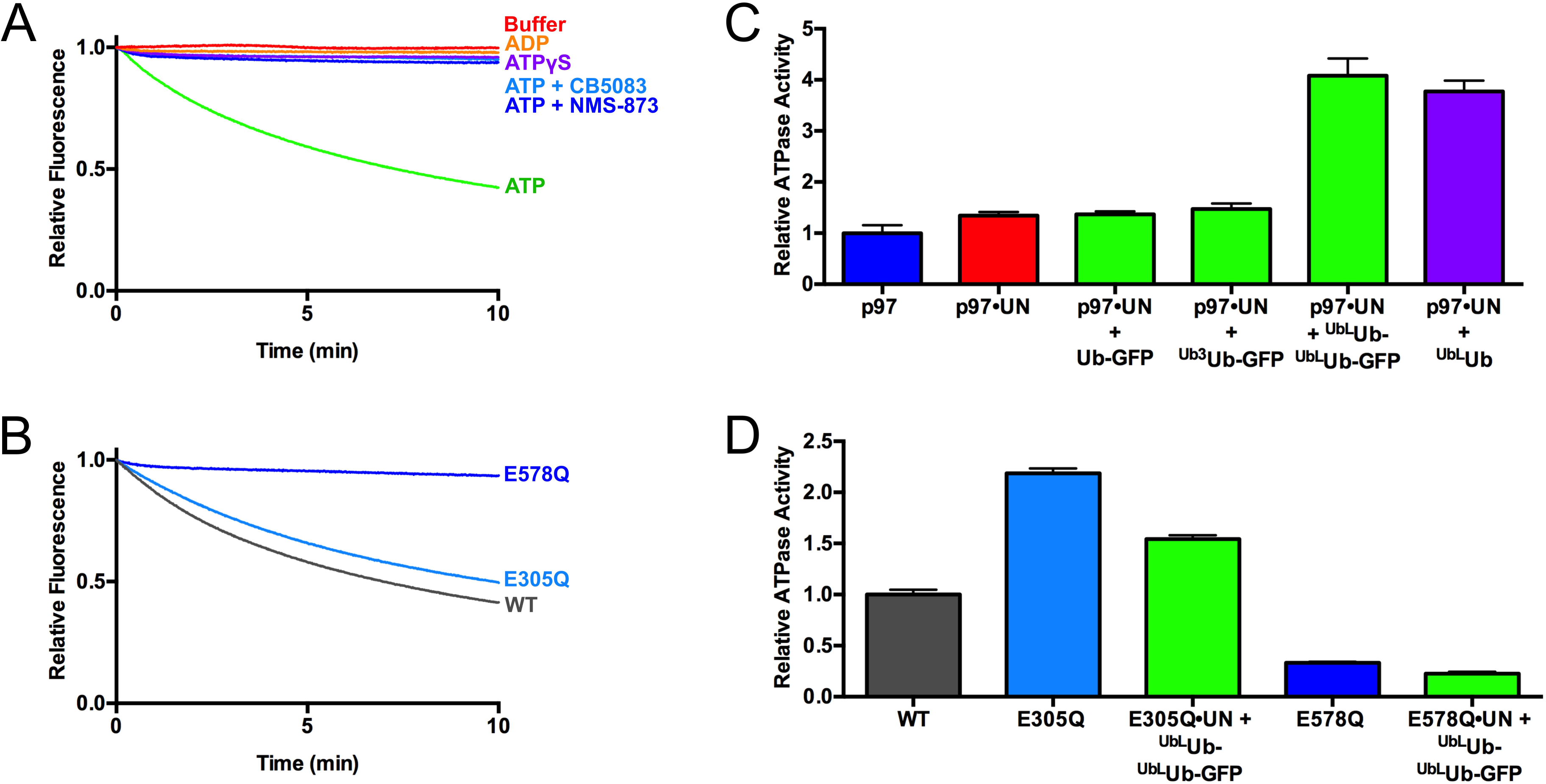
ATPase activity of p97 is critical for and stimulated by substrate unfolding. (*A*) Fluorescence traces of ^UbL^Ub-^UbL^Ub-GFP in the presence of p97, UN, GroEL, and various nucleotides and p97 inhibitors. Unfolding was observed only in the presence of ATP. Representative traces shown, n≥2. (*B*) Unfolding of ^UbL^Ub-^UbL^Ub-GFP by p97 ATPase mutants. p97-E305Q and p97-E578Q are deficient in D1 and D2 ATPase activity, respectively. p97-E305Q was able to unfold substrate whereas p97-E578Q was not. Representative traces shown, n≥3. (*C*) Substrate stimulates ATPase activity of p97 when UN is present. Both unanchored Ub chains and those linked to Ub-GFP yielded equivalent stimulation, while Ub-GFP and ^Ub^3^^Ub-GFP did not stimulate. ATPase activity was measured with BioMol Green as described in Methods and was normalized to basal WT p97 activity. Error bars represent S.D. with n=4. (*D*) Effect of substrate plus UN on ATPase activity of D1 and D2 domain ATPase mutants. Addition of ^UbL^Ub-^UbL^Ub-GFP plus UN slightly decreased ATPase activities of both D1 mutant E305Q and D2 mutant E578Q. Error bars represent S.D. with n=4.

**Table 1.** Rates and extents of unfolding of ^Ub^^L^Ub-^UbL^Ub-GFP by p97 mutants. The rates and plateaus were calculated by fitting data to a single exponential decay, with the plateau representing the percent of fluorescence remaining at the end of the reaction. Unpaired t-tests comparing WT rates to those of p97-E305Q and p97-A232E yielded p-values of <0.0001 in both cases, indicating statistically significant differences. Sample size represents number of technical replicates, and values are shown ± S.D.

| **Mutant** | **Rate (s<sup>-1</sup>)** | **Plateau (%)** | **n** |
| --- | --- | --- | --- |
| WT | 0.195 ± 0.007 | 33.3 ± 0.8 | 6 |
| E305Q | 0.154 ± 0.005 | 36.5 ± 0.8 | 3 |
| E578Q | N.D. | N.D. | 3 |
| A232E | 0.274 ± 0.007 | 25.3 ± 0.4 | 3 |

Some adaptors modulate p97 ATPase activity (38), so we examined the effects of substrate processing on the hydrolysis of ATP by p97 and p97•UN. Addition of long unanchored K48-linked ubiquitin chains (^UbL^Ub), Ub-GFP, ^Ub^3^^Ub-GFP or ^UbL^Ub-^UbL^Ub-GFP did not alter the ATPase activity of p97 (Fig. S7*A*). Whereas UN did not significantly affect the ATPase activity of p97, the further addition of ^UbL^Ub-^UbL^Ub-GFP stimulated ATP hydrolysis by ∼4-fold, whereas Ub-GFP and ^Ub3^Ub-GFP had no effect (Fig. 4C). The latter result is consistent with the inability of p97•UN to unfold Ub-GFP or ^Ub^3^^Ub-GFP. Neither NSFL1C nor UBXN7 supported substrate-triggered acceleration of p97 ATP hydrolysis (Fig. S7*B*). Long, unanchored Ub chains also stimulated p97 ATPase activity to a similar degree as ^UbL^Ub-^UbL^Ub-GFP, suggesting that p97 interaction with Ub chains and not the GFP substrate itself was both necessary and sufficient for the observed acceleration. In agreement with the unfolding results, the residual ATPase activity of p97-E578Q was unaffected by substrate plus UN (Fig. 4D). However, p97-E305Q, which was able to unfold substrate, was also not stimulated by substrate, even showing a slight decrease in ATPase activity (Fig. 4D). The E305Q mutant also showed higher basal ATPase activity than WT, as has been seen before in steady-state experiments (20). These results suggest that there is a high degree of crosstalk between ATPase activity in D1 and D2, and whereas ATP hydrolysis in D2 is the driving force for unfolding, D1 activity is also needed for substrate-induced ATPase acceleration.

### UN recruits substrate to p97

The tight correlation between the competence of a substrate to be unfolded and its ability to accelerate ATP hydrolysis suggests that binding of substrate to p97 may stimulate ATPase activity, leading to substrate unfolding. To further probe this hypothesis, we evaluated binding of substrate to p97. Immunoprecipitation of p97 showed that it bound ^UbL^Ub-^UbL^Ub-GFP substrate in the presence but not in the absence of UN (Fig. 5A, lanes 4 and 5). Reciprocal immunoprecipitation of GFP confirmed that substrate bound UN in the absence of p97 but did not bind p97 in the absence of UN (Fig. 5B). Furthermore, the interaction between p97 and UN appeared to be stabilized by substrate binding (Fig. 5A, lanes 3 and 5). We also analyzed substrate dependence of GroEL association with p97, but observed high background binding to our beads even in the absence of antibody (lane 1). The binding signal was increased in lanes 6 and 7, but this could be due to p97, which in our experience is prone to exhibit non-specific binding.

**Fig. 5.**
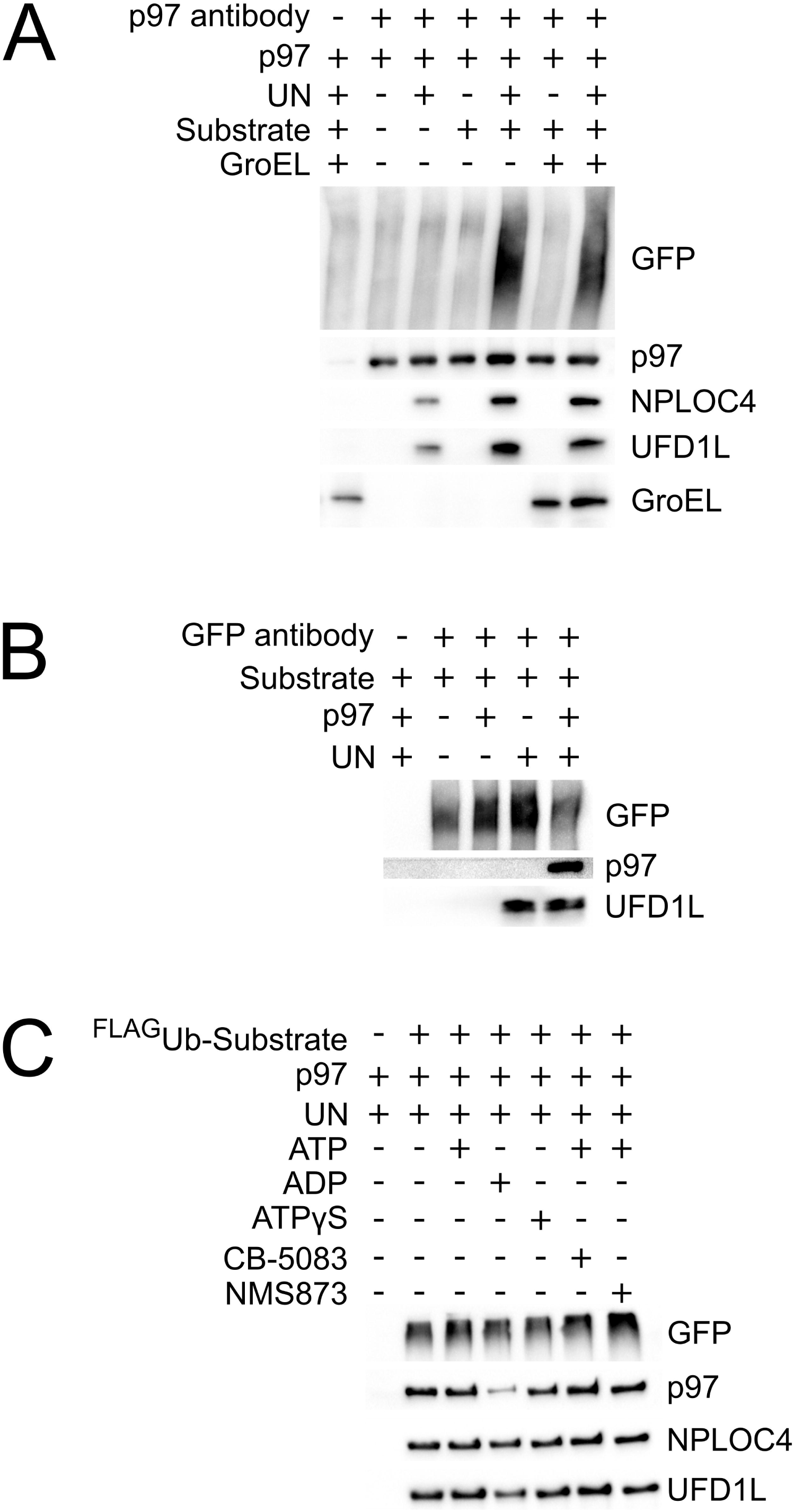
UN recruits ubiquitylated substrate to p97. (*A*) Association of substrate with p97 depends on UN. Reactions containing 100 nM p97, 200 nM UN, 200 nM ^UbL^Ub-^UbL^Ub-GFP and/or 200 nM GroEL were immunoprecipitated using an anti-p97 antibody and assessed by western blot. Substrate was pulled down with p97 only in the presence of UN, and substrate enhanced the binding of UN to p97. Effects of substrate and UN on binding of GroEL to p97 could not be determined due to high background binding of GroEL to beads, antibody and/or p97. (*B*) UN binds directly to substrate and links it to p97. Samples prepared as in (*A*) were immunoprecipitated with an anti-GFP antibody. Substrate pulled down UN but only bound p97 in the presence of UN. (*C*) Bound nucleotide has only a modest effect on formation of a p97•UN•substrate ternary complex. Samples prepared as in (*A*) with various nucleotides and/or p97 inhibitors were immunoprecipitated by anti-FLAG resin, which bound the FLAG-tagged Ub on substrate. Binding of p97, but not UN, was reduced only in the ADP state.

Whereas the unfolding of substrate was highly dependent on ATP hydrolysis, substrate interaction with p97 was not. Immunoprecipitation of ^FLAG-UbL^Ub-^FLAG-UbL^Ub-GFP pulled down equal amounts of p97 in the absence of added nucleotide and in the presence of ATP, ATPγS, ATP plus NMS-873, and ATP plus CB-5083 (Fig. 5C). Therefore, p97 does not have to be actively remodeling substrate in order to effectively form a tight complex. Loss of binding of p97, but not UN, was seen only with added ADP, which could be due to the large conformational changes observed for p97 in its ADP-bound state (18) (Fig. 5C). These results are consistent with previous studies on Ub chain association with p97 and p97•UN (48, 82).

### IBMPFD mutant p97-A232E has enhanced unfoldase activity

The autosomal dominant human syndrome IBMPFD is caused by mutations that cluster at the interface of the N and D1 domains of p97. Of the different IBMPFD mutations that have been identified in p97, A232E exhibits the most severe phenotype in terms of age of onset and penetrance. In addition, p97-A232E consistently shows higher basal D2 ATPase rates than wild-type enzyme (20, 63, 65). The availability of an assay that directly measures ATP-dependent unfolding by p97 allowed us to distinguish between two alternative interpretations of the significance of the enhanced ATPase activity of the A232E mutant protein. On the one hand, the enhanced activity may reflect a true gain-of-function wherein p97-A232E is either a more powerful or faster motor. On the other hand, ATPase activity may increase because of decoupling of the D2 “motor” from the substrate “load”, analogous to pushing in the clutch when an engine is revving in low gear. If the former is more accurate we would expect to see increased unfolding by the mutant protein. Conversely, if the latter is correct, we would expect to see reduced unfolding. The result of this experiment was unambiguous: p97-A232E exhibited accelerated unfolding of GFP under the single-turnover conditions used in the assay (Fig. 6A, Table 1). This effect is reproducible, because in two independent sets of preparations, p97-A232E displayed faster unfolding than WT p97 (Fig. 6A and Fig. 6C). The substrate-induced ATPase acceleration observed for WT p97 was also observed with p97-A232E (Fig. 6B). The mutant protein displays an increased basal rate of ATP hydrolysis and a higher rate in the presence of substrate compared to WT.

**Fig. 6.**
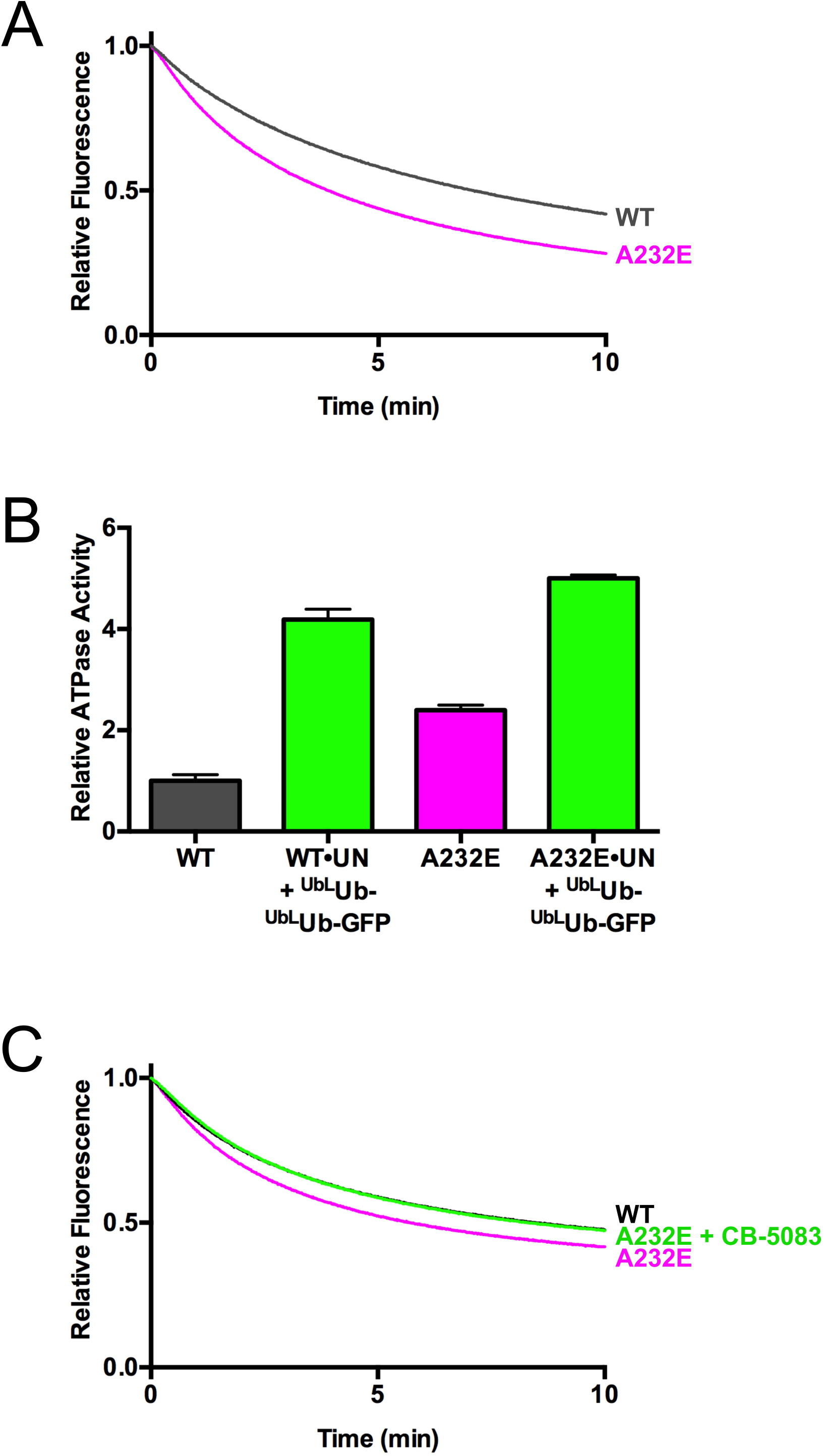
IMBPFD mutant p97-A232E is a better unfoldase. (*A*) In the presence of UN and GroEL, p97-A232E catalyzed the loss of fluorescence of ^UbL^Ub-^UbL^Ub-GFP faster than wild-type, suggesting it acts as an improved unfoldase. Rates are listed in Table 1 and the difference between WT and p97-A232E is statistically significant with p<0.0001. Representative traces shown, n ≥ 3. (*B*) p97-A232E shows accelerated ATPase rates in the presence of UN and substrate. The IBMPFD mutant also had both a higher baseline ATPase rate than WT (unpaired t-test, p<0.0001) and a higher ATPase activity in the presence of UN and substrate compared to WT (unpaired t-test, p=0.0003). ATPase activity was measured with BioMol Green as described in Methods was normalized to basal WT p97 activity. Error bars represent S.D. with n=4. (*C*) Addition of a 37.5 nM CB-5083 restores p97-A232E unfoldase activity to WT levels. Representative traces shown, n=2, and the difference between WT and p97-A232E rates is statistically significant (unpaired t-test, p=0.02). Independent preparations of proteins were used for panels (*A*) and (*C*).

We identified a reversible, ATP-competitive p97 ATPase inhibitor that was recently perfected to yield the clinical candidate CB-5083 (81), which is in human phase 1 clinical trials for treatment of cancer. If the basis of IBMPFD pathology is due to enhanced unfoldase activity of p97, we reasoned that CB-5083 could potentially be explored as a therapy for IBMPFD. To test the feasibility of this idea, we performed unfolding reactions with p97-A232E at several different concentrations of CB-5083. Remarkably, when added at 37.5 nM, or 1 molecule per 2 p97 hexamers, CB-5083 normalized the unfolding rate of p97-A232E to match that of WT (Fig. 6C).

## Discussion

P97/VCP is implicated in a broad range of cellular processes including membrane fusion, protein trafficking, and ubiquitin-dependent proteolysis (1). It is thought that the core biochemical activity of p97 that enables its diverse biological functions is its ability to act as a ‘segregase’ that segregates polypeptides from binding partners in multisubunit complexes, or from large macromolecular structures including ribosomes, membranes, or chromatin. Although the mechanism by which p97 exerts segregase activity is not known, the most economical hypothesis is that it grabs onto the polypeptide to be segregated and commences to unfold it. However, despite the appeal of this unifying hypothesis, the ability of wild type p97 to harvest the energy of ATP hydrolysis to unfold a polypeptide has never been directly demonstrated. We show here for the first time, and in a well-defined system, that wild type p97 can at least partially unfold a protein. In contrast to prior studies that employed a doubly-mutated p97 acting upon an unmodified protein in the absence of any adaptor (83, 84), the unfolding we observe exhibits dependencies predicted from prior genetic and biochemical studies of p97, including dependence on ATP hydrolysis by the D2 ATPase domain of p97, the heterodimeric adaptor NPLOC4•UFD1L, and conjugation of an ubiquitin chain to the substrate that is unfolded (1, 24, 25).

Despite the close alignment of in vitro dependencies reported here with known in vivo requirements for p97 action, there are two caveats worth noting. First, we do not know whether the loss of GFP fluorescence is due to complete unfolding or local unfolding. Second, our Ub-GFP model substrate is not a native substrate of p97. However, p97 activity in the UFD pathway is absolutely required for its degradation in vivo in yeast, *Drosophila*, and human cells (19, 47, 70, 72), and the E2–E3 chimera employed to ubiquitylate Ub-GFP was derived from enzymes that function directly upstream of p97 in the ERAD pathway (73). Thus, we believe the reaction reported here represents the essence of p97’s biochemical activity that underlies its physiological functions.

Processing of our Ub-GFP substrate by p97 is dependent on the conjugation of multiple ubiquitins connected by K48 linkages, which is consistent with reports of K48-linked Ub chain binding by p97 and UFD1L and linkage-nonspecific chain binding by NPLOC4 (25, 48, 49, 82). One of our most interesting findings is that we observed enhanced substrate unfolding when the substrate contained a branch point enabling the formation of multiple K48-linked ubiquitin chains. Our substrate contains a branch at the base of the ubiquitin conjugate, which allows for two K48-linked chains to be built on a single site of substrate modification. Because there are multiple Ub binding sites on the p97•UN complex, the branched chain on our ^UbL^Ub-^UbL^Ub-GFP substrate may retard, via enhanced avidity, its dissociation prior to unfolding. Branched Ub chains can enhance substrate degradation (85) and have been associated with p97 previously. Ubiquitin chains with K11 linkages are associated with multiple p97 adaptor proteins (37) and are implicated in ERAD (86). Substrates modified with K11 and K11-K48 branched chains associate with p97•UN and its *Drosophila* ortholog (87, 88), and p97 binds K11-K48 branched chains better than either type of homotypic chain (85). Furthermore, both K29 and K48 linkages are formed on UFD pathway substrates in vivo and in vitro using purified UFD pathway enzymes (47, 89). Whereas our data with linear ubiquitin fusions suggest branched and/or multiple ubiquitin chains may be important for interaction of substrates with p97•UN, further work with physiological linkages in vitro and in vivo is needed to determine the exact requirements for ubquitylated p97 substrates.

The second requirement for p97-catalyzed unfolding is the heterodimeric adaptor UN. In agreement with prior work, UN bridges the interaction of substrates with p97 (25, 48). However, p97 also associates with other adaptor proteins, some of which (e.g. UBXN7) can co-bind with UN and others (e.g. NSFL1C) which bind in a mutually exclusive manner (36, 37, 46, 90, 91). Although there are examples of processes that require both UN and an additional adaptor (42, 92), in general it is unclear whether the different adaptors work in different pathways, or work sequentially, together, or in opposition to one another in the same pathway. Mutually exclusive adaptors like NSFL1C bind very differently from UN yet also promote protein segregation (41, 51, 93). However, neither NSFL1C nor UBXN7 was able to replace UN in our unfolding assay, suggesting they promote substrate processing in a different way or work on different substrates. Our assay provides the first platform for further mechanistic exploration of the functional relationship between different adaptor proteins.

The final requirement for unfolding of GFP is ATP hydrolysis by the D2 ATPase of p97. Our results are in direct contrast with a previous study which showed that p97-dependent unfolding is inhibited by ATP (94), but are consistent with overwhelming in vitro and in vivo data (1, 23, 25, 34). Our experiments with p97 mutants and the D2-specific ATP-competitive inhibitor CB-5083 demonstrate the importance of D2 ATPase activity over that of D1, which confirms prior observations (19–25). We observed very little effect of the D1 Walker B mutant on substrate processing, leaving the role of D1 ATPase activity in p97 function unclear (1). Not only is the ATPase activity key to substrate processing but we also observe its stimulation by Ub chain or substrate binding. This stimulation was abolished in the D1 Walker B mutant. Nevertheless, this mutant unfolded GFP at near-WT rates, suggesting that substrate stimulation of ATP hydrolysis might not be essential, at least for some substrates. ATPase acceleration in the presence of the cytoplasmic fragment of the ERAD substrate Sytl has been previously reported, but unlike the stimulation reported here, it was not dependent upon UN or ubiquitylation of the substrate (34), both of which are thought to be requirements for ERAD. p97 can interact both with a substrate and the Ub chain appended to the substrate (25, 95), suggesting that stimulation of p97 ATPase and perhaps substrate processing can be regulated through multiple binding interfaces. Though we are unable to draw any conclusions about the mechanism of unfolding by p97, studies on the archaebacterial homolog VAT support a model whereby substrate is translocated through the central channel (14, 15), suggesting that adaptor and substrate engagement must expand p97’s narrow pore (16, 18) to accommodate the threading of an unfolded polypeptide chain.

Mutations in p97 cause the autosomal dominant human disease IBMPFD (53). Prior work has led to conflicting proposals regarding the underlying basis for pathogenesis in IBMPFD. On the one hand, some studies have emphasized a reduction in specific biological functions of the mutant p97, including its roles in endosomal trafficking, autophagy, and elimination of leaky lysosomes (56, 8, 60). Defects in these processes have been linked to reduced binding of mutant p97 to UBXD1 (8). On the other hand, IBMPFD mutant proteins hydrolyze ATP at a faster rate (20, 63–65). Whereas it has been speculated that the increase in ATP hydrolysis might be due to an uncoupling of substrate binding via the N domain to mechanochemical transduction in the D2 domain (54, 68), studies in *Drosophila* point to an increase in function for the mutant p97 (61, 67, 69).

Furthermore, the overexpression of WT p97 enhanced IBMPFD mutant phenotypes while inactivation of one copy of WT p97 suppressed mutant phenotypes, which is consistent with a gain-of-function mutation (96). The availability of a biochemical assay that directly measures substrate unfolding, which we propose is the core biochemical activity underlying p97’s myriad biological functions, allowed us to investigate this issue. We focused on the mutant with the most severe disease phenotype, p97-A232E, and observed a modest but reproducible increase in the rate of substrate unfolding that was re-normalized by addition of an ATP-competitive inhibitor, implying that the underlying defect in this disease may be due, in part, to a true gain-of-function rather than the prevailing loss-of-function hypothesis. Our results call to mind previous work to engineer Hsp104, in which point mutations greatly augmented its ATPase and unfoldase activity while altering intersubunit communication (97). It is interesting that the increased ATPase activity of p97-E305Q did not cause a similar increase in unfolding rate, suggesting that the rate of ATP hydrolysis in D2 does not by itself determine the rate of unfolding. Our experiments were carried out with pure WT and pure mutant p97, so it will be of interest to study populations of mixed hexamers of A232E and other IBMPFD mutants, to better recapitulate the situation that pertains in vivo. Whereas much clearly remains to be done to further investigate this gain-of-function model, if our proposal is correct it implies that the clinical-grade p97 inhibitor (81) that renormalized the activity of the mutant protein may be useful for therapy of IBMPFD and ALS cases that arise from mutation of p97.

## Methods

### Protein and expression purification

A description of the construction, expression and purification of novel reagents is described below. A description of previously published proteins used in this study can be found in Table S1.

### gp78RING-Ube2g2 chimera construction and purification

A 72-residue sequence of the E3 ubiquitin protein ligase gp78/AMFR (residues 322 to 393), containing the RING domain, was fused to the N-terminus of the ubiquitin-conjugating enzyme Ube2g2 with a linker sequence of GTGSH. cDNA encoding this fusion protein was inserted into the bacterial expression plasmid p28a-TEV vector to encode a polyHis-tagged protein. Protein was expressed in BL21(DE3) at 37 °C with 0.4 mM IPTG for 4 hours and was purified on Ni-NTA resin and a Superdex 75 column before being cleaved with TEV protease overnight at 4 °C. Cleaved protein was then bound to a MonoQ ion exchange column, eluted with a NaCl gradient (0.05 – 0.5 M), concentrated with centrifugal filter units, and flash frozen.

### Ub^G^76^V^GFP fusion construction

The coding sequence for Ub^G^76^V^GFP in the EGFP-N1 vector (RDB# 1832) (19) was PCR amplified and inserted into pET28a using NdeI and NotI sites to produce His_6_-Ub^G^76^V^-GFP (RDB# 3006). His_6_-Ub^G^76^V^- Ub^G^76^V^ GFP (RDB# 3344) and His_6_-Ub^G^76^V^-Ub^G^76^V^-Ub^G^76^V^ -GFP (RDB# 3345) were created by ligating Ub_2_^G76V^ and Ub_3_^G76V^, PCR amplified from a synthetic Ub_4_^G76V^ sequence (RDB# 2406), into His-Ub^G^76^V^-GFP cut with NdeI and HindIII. For simplification, the G76V notation has been left out of subsequent mentions of these constructs.

### ^Ub3^Ub-GFP synthesis and purification

A plasmid for bacterial expression of Ub-K48R (RDB# 3348) was made from that of Ub (RDB# 2805) by site-directed mutagenesis, and the protein was expressed as previously described (76). Pure K48-linked Ub_3_ chains carrying a K48R mutation in the distal Ub were enzymatically synthesized and purified as described previously (76, 77). To form ^Ub^3^^Ub-GFP, 0.5 μM Ube1 (E1), 5 μM gp78RING-Ube2g2, 2.5 μM Ub-GFP, and 5 μM Ub_3_ were incubated in 20 mM Hepes pH 7.4, 5 mM ATP, and 5 mM MgCl_2_ overnight at 37 °C. The ^Ub^3^^Ub-GFP was then purified on Ni-NTA resin and a Superdex 200 gel filtration column before being concentrated with centrifugal filter units and flash frozen.

### Polyubiquitylated substrate synthesis and purification

Reactions were carried out with final concentrations of 10 μM Ub-GFP fusion protein, 1 μM E1, 20 μM gp78RING-Ube2g2, and 400 μM ubiquitin in 20 mM Hepes pH 7.4, 10 mM ATP, and 10 mM MgCl_2_ at 37 °C overnight. Ubiquitin was added progressively in small amounts over the first eight hours of the reaction. For FLAG-tagged substrate, 40 μM FLAG-Ubiquitin (Boston Biochem, Cambridge, MA, USA) was added in the first two hours. To purify ubiquitylated GFP from free ubiquitin chains, the reaction mixture was incubated with Ni-NTA resin, eluted with 300 mM imidazole, and run over a Superose 6 size exclusion column in 20 mM Hepes, pH 7.4, 250 mM KCl, 1 mM MgCl_2_, 1 mM TCEP and 5% glycerol. Fractions were pooled into long, medium, and short chain length samples, concentrated with centrifugal filter units, and flash frozen.

### ATPase assays

The ATPase assay protocol was modified from previously published methods (40). In an untreated microplate (#655101, Greiner Bio-One, Kremsmünster, Austria), 40 μL solutions containing 30 nM p97 hexamer, 150 nM adaptor, and/or 150 nM substrate in ATPase assay buffer (25 mM Hepes pH 7.4, 100 mM KCl, 3 mM MgCl_2_, 1 mM TCEP, 0.1 mg/mL ovalbumin) were preincubated at 37 °C for 10 minutes. To this, 10 μL of a 1 mM ATP solution was added, and the reaction was incubated at 37 °C for 5 or 10 minutes. After cooling on ice for 30 seconds, 50 μL of BIOMOL Green reagent (Enzo Life Sciences, Farmingdale, NY, USA) was added. Solutions were developed at room temperature for 30 minutes before being read at 600 nm. The amount of inorganic phosphate in each sample was calculated from a standard curve, and relative ATPase activity for a sample was calculated by normalizing its measurement to that of samples of WT p97 alone.

### Fluorescence unfolding assays

Unless specified, all assays were carried out at 37 °C. Samples contained 25 nM GFP substrate, 75 nM p97 hexamer, 150 nM adaptor (UN, NSFL1C or UBXN7), and/or 250 nM GroEL trap in unfolding assay buffer (25 mM Hepes pH 7.4, 100 mM KCl, 5 mM MgCl_2_, 1 mM TCEP, 2 mM ATP). Control experiments indicated that the levels of p97, adaptor, and GroEL used were at or near saturation (Figure 2—figure supplement 4). Other nucleotides and p97 inhibitors were present at 2 mM and 10 μM, respectively, when indicated. Kinetic experiments were carried out on a Fluoro-Log 3 (Horiba Jobin Yvon, Edison, NJ, USA) with excitation at 488 nm and emission at 509 nm. Relative fluorescence was calculated by normalizing the fluorescence signal to that at time zero. Unfolding rates were calculated by fitting curves to an exponential decay model in Prism (GraphPad, San Diego, CA, USA).

### Binding assays

Antibodies used for immunoprecipitation and western blotting are listed in Table S2. Samples containing 100 nM p97, 200 nM UN, 200 nM GFP substrate, and/or 200 nM GroEL trap in binding assay buffer (25 mM Hepes pH 7.4, 100 mM KCl, 3 mM MgCl_2_, 1 mM TCEP) in a volume of 200 μL were preincubated at 37 °C for 15 minutes. Triton X-100 interfered with binding of substrate to GroEL trap, so 0.01% Triton X-100 was included in all buffers only in reactions that did not contain GroEL. Nucleotides and inhibitors were present at 2 mM and 10 μM, respectively, where indicated. For IPs using Protein G magnetic beads (Bio-Rad Laboratories, Hercules, CA, USA), protein mixtures were incubated with 1 μL of antibody for 15 minutes at 37 °C prior to a 1 hour incubation with 25 μL of beads at room temperature. For FLAG IPs, reactions were incubated with 25 μL of anti-FLAG resin (Sigma-Aldrich, St. Louis, MO, USA) for 15 minutes at room temperature. Following incubation, all beads were washed 3 times with 750 μL assay buffer before being boiled in 50 μL 2X SDS-PAGE loading dye. Samples were then analyzed by western blot.

## Acknowledgements

We thank Willem den Besten, David Sherman, and Jing Li for assistance with cloning and protein purification, Arthur Horwich for the GroEL expression plasmid and antibody, Rati Verma and David Sherman for comments on the manuscript, and the entire Deshaies lab for helpful discussion. We also thank Tom Rapoport for communicating results prior to publication. Fluorescence measurements were carried out in the Beckman Institute Laser Resource Center and the Caltech Biophysical Facility. R.J.D. is an Investigator of the HHMI and this work was supported by HHMI.

## Competing interests

RJD: RJD is also a founder, consultant, member of the SAB, and shareholder of Cleave Biosciences, which is developing CB-5083 for therapy of cancer. The other authors declare that no competing interests exist.

## Author contributions

EEB designed, performed, and interpreted all experiments related to p97, and drafted and edited the manuscript; KCO designed and characterized gp87RING- Ube2g2 and edited the manuscript; VC assisted in the design and characterization of gp78RING-Ube2g2 chimera and edited the manuscript; RJD assisted in the design and interpretation of all experiments related to p97, and drafted and edited the manuscript.

